# SoyOmics: A deeply integrated database on soybean multi-omics

**DOI:** 10.1101/2023.02.05.527102

**Authors:** Yucheng Liu, Yang Zhang, Xiaonan Liu, Yanting Shen, Dongmei Tian, Xiaoyue Yang, Shulin Liu, Lingbin Ni, Zhang Zhang, Shuhui Song, Zhixi Tian

## Abstract

As one of the most important crops to supply majority plant oil and protein for the whole world, soybean is facing an increasing global demand. Up to now, vast multi-omics data of soybean were generated, thereby providing valuable resources for functional study and molecular breeding. Nevertheless, it is tremendously challenging for researchers to deal with these big multi-omics data, particularly considering the unprecedented rate of data growth. Therefore, we collect the reported high-quality omics, including assembly genomes, graph pan-genome, resequencing and phenotypic data of representative germplasms, transcriptomic and epigenomic data from different tissues, organs and accessions, and construct an integrated soybean multi-omics database, named SoyOmics (https://ngdc.cncb.ac.cn/soyomics). By equipping with multiple analysis modules and toolkits, SoyOmics is of great utility to facilitate the global scientific community to fully use these big omics datasets for a wide range of soybean studies from fundamental functional investigation to molecular breeding.

## Dear Editor

As one of the most important crops to supply majority plant oil and protein for the whole world, soybean is facing an increasing global demand (Ray et al., 2013). The reference genome of accession “Williams82” opened the gate of genomics research in soybean (Schmutz et al., 2010). After that, vast multi-omics data were generated, thereby providing valuable resources for functional study and molecular breeding (Zhang et al., 2022). Nevertheless, it is tremendously challenging for researchers to deal with these big multi-omics data, particularly considering the unprecedented rate of data growth (Yang et al., 2021; Cao et al., 2022). Thus, constructing an integrated multiomics database for soybean that provides a one-stop solution for big data mining with friendly interactivity is highly desired.

Here, we collect the reported high-quality omics data (Figure 1A), including assembly genomes, graph pan-genome, resequencing and phenotypic data of representative germplasms (Zhou et al., 2015; Fang et al., 2017; Liu et al., 2020), transcriptomic (Shen et al., 2014; Shen et al., 2019) and epigenomic (Shen et al., 2018) data from different tissues, organs and accessions, and construct an integrated soybean multi-omics database, named SoyOmics (https://ngdc.cncb.ac.cn/soyomics). By equipping with multiple analysis modules and toolkits, SoyOmics is of great utility to facilitate the global scientific community to fully use these big omics datasets for a wide range of soybean studies from fundamental functional investigation to molecular breeding.

**Figure 1.**
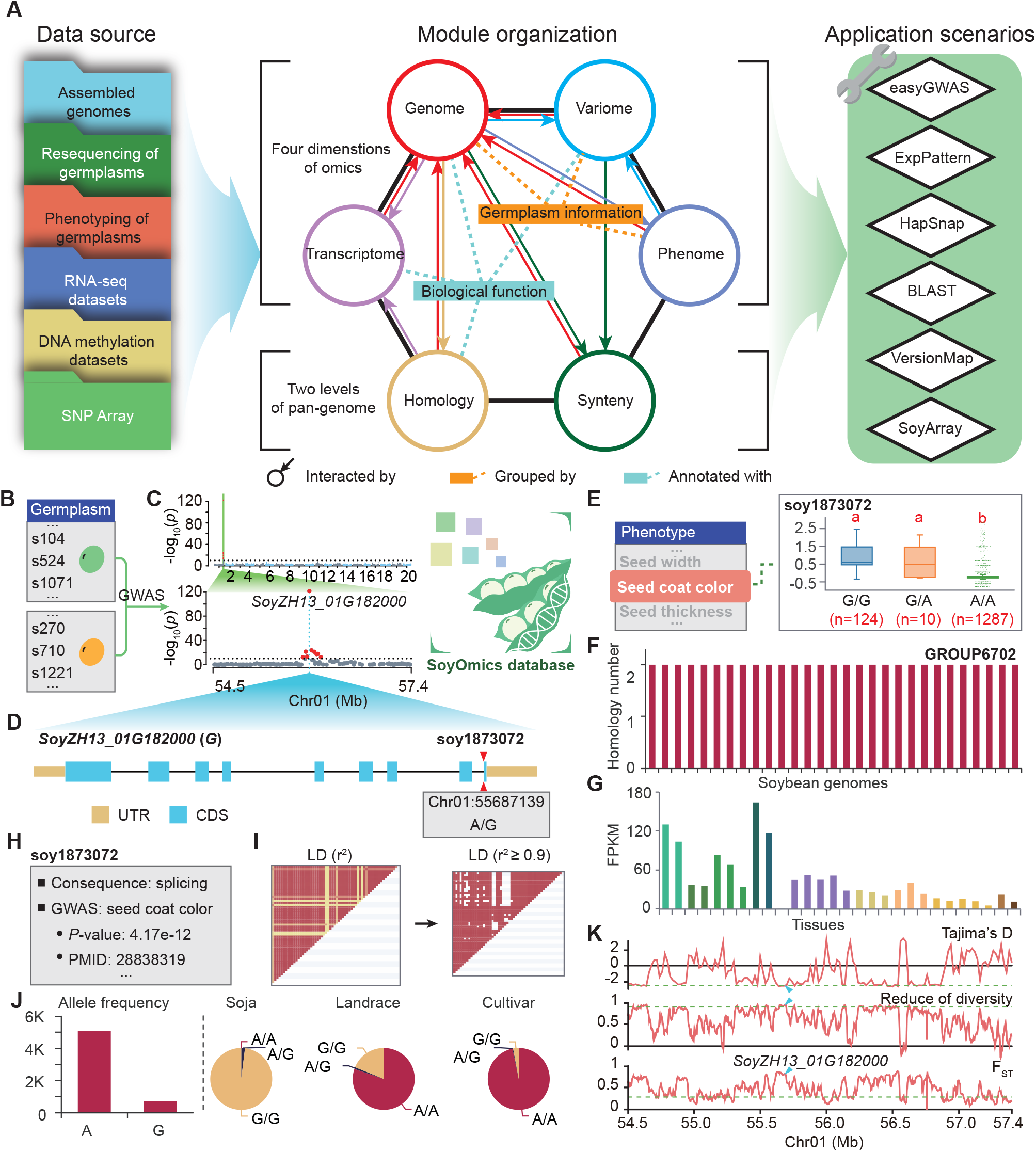
Overview of SoyOmics and its practice for data mining. (A) Framework of SoyOmics, including data resource, module organization and application scenarios. (B) Germplasms grouped by seed coat color. (C) GWAS analysis with seed coat color. Threshold is set by *P* = 1e-10. (D) Gene structure and variation list of *SoyZH13_01G182000*. (E) Seed coat color grouped by genotypes of soy1873072. Multiple comparison is conducted by Student-Newman-Keuls (SNK) test, with significant level equal to 0.01. (F) Homologous gene number of *SoyZH13_01G182000* in 29 soybean genomes. (G) Expression of *SoyZH13_01G182000* in 28 tissues. (H) Change consequence on mRNA and present knowledge of variation soy1873072. (I) Linkage disequilibrium heat map around soy1873072. The left chart shows heat map of all variation pairs in the block, and the right shows heat map of variation pairs with r2 ≥ 0.9. (J) Genotype frequency distribution of *G* gene in *Soja*, landrace and cultivar. (K) Selective test signal of genomic region Chr01:54.5-57.4 Mb. Green dash line means the significant threshold. Blue triangle shows the location of *SoyZH13_01G182000*.

## An overview of SoyOmics

By integrating different multi-omics data, we develop six highly interactive basic modules in SoyOmics: Genome, Variome, Transcriptome, Phenome, Homology, and Synteny (Figure 1A). The Genome module embodies the information of 2,898 soybean germplasms and 27 *de novo* assembled genomes, providing users with open access to basic information of sequenced germplasms, individual genomes and genes (Supplemental Figure 1). The Variome module organizes approximately 38 million SNPs and INDELs of the 2,898 soybean accessions, facilitating users to check the variation information and whole genome selective signals for any germplasm of interest (Supplemental Figure 2). The Transcriptome module contains two datasets of gene expression: one is from 27 tissues at different developmental stages from Williams82 and ZH13 accessions, respectively, and the other is from 9 tissues at different developmental stages from each of the 26 accessions used for pan-genome analysis. In this module, users can obtain gene expression profiles and gene orthologous information by specifying gene ID or functional description (Supplemental Figure 3). The Phenome module collects approximately 27 thousand records of 115 phenotypes with terms defined as controlled vocabularies that fall into 5 classes (including morphology, growth and development, biochemistry, biotic stress, and vigor) as well as 17 subclasses (Supplemental Figure 4). The Homology module displays the soybean pan-genome by characterizing 57,480 homologous gene groups. Users can specify any gene ID, homologous group ID or gene functional description to retrieve the homologous group of interest (Supplemental Figure 5). The Synteny module deposits approximately 550 thousand large-scale structural variations (SVs) in the pan-genome, in which users can visualize and download the SVs and synteny blocks by setting a specific genomic region. Furthermore, the graph pan-genome is embedded and a SequenceTubeMap web service (https://github.com/vgteam/sequenceTubeMap) is deployed for visualization of pan-genome threads (or haplotypes) according to nodes made up by SVs (Supplemental Figure 6).

In addition, SoyOmics is designed to provide user-friendly search bar in each module and to cover as much as more possible substances. According to the searching category and inputting context, it features powerful search engine to provide comprehensive associated results with friendly links from one module to other modules (Supplemental Figure 7).

## Application toolkits

In addition to the six modules, we design several commonly easy-to-use toolkits, including *easyGWAS, ExpPattern, HapSnap, BLAST, VersionMap*, and *SoyArray* (Figure 1A). The *easyGWAS* is a tool for quick-start genome-wide association study (GWAS) analysis, providing friendly interface for parameters setting and algorithms selection and offering multiple high-quality outputs including Manhattan plot, QQ-plot and text result (Supplemental Figure 8). The *ExpPattern* is for conducting expression pattern analysis for a gene list against soybean tissues. It can generate expression heatmap, with options of whether to execute clustering or not (Supplemental Figure 9). Besides, the tspex is incorporated in the *ExpPattern* for advice of gene’s tissue-specificity (Camargo et al., 2020). The *HapSnap* is designed for haplotype analysis for a genomic region. Users can refine the variations via selection of variation type and quality control. The output includes haplotype frequency, haplotype vs. genotype, and linkage disequilibrium (Supplemental Figure 10). The *VersionMap* is capable to convert the genomic region between ZH13 (v2) and other *de novo* genomes of soybean, or gene ID between Williams82 (v2) and ZH13 (v2) (Supplemental Figure 11).

We also develop a toolkit named *SoyArray* by embedding the information of GenoBaits soybean array (Liu et al., 2022), in which users can search and download the marker information they are interested in. We also afford a function in the *SoyArray* to compare divergent sites between two germplasms based the makers from GenoBaits soybean array, which is helpful for parents’ picking in genetic or breeding study (Supplemental Figure 12).

## Data mining using SoyOmics

As SoyOmics integrates a wide variety of soybean multi-omics data, it can be used for deep mining ranging from fundamental research to molecular breeding. Here we take a previously reported seed coat color causal gene, *G* (Wang et al., 2018), as an example. In SoyOmics, we can group germplasms by green or yellow seed coat colors (Figure 1B). According to the phenotype data, we can easily conduct GWAS using the *easyGWAS* toolkit, and then identify a significant association signal that is located in the *G* gene, *SoyZH13_01G182000* (Figure 1C and 1D). According to the interested association genetic variant, users can get phenotype variations among different genotypes, such as the seed coat color (Figure 1E). By searching the candidate gene *SoyZH13_01G182000* from different modules, users can obtain a wealth of gene information including basic summary, functional annotation, homology in 29 soybean genomes and expression pattern in 28 tissues (Figure 1D, 1F, and 1G). Furthermore, users can also investigate functional annotations for any variant of interest (Figure 1H), linkage disequilibrium around the association genetic variant (Figure 1I), allele frequency in different populations (Figure 1J), and selection sweeps for the association regions by three different test methods (Figure 1K). Notably, the majority of charts generated in SoyOmics can be directly downloaded and edited.

In summary, SoyOmics features comprehensive integration of multi-omics datasets and provides user-friendly interfaces for soybean study. Undoubtedly, soybean omics data are generated at increasing scales and rates, including resequencing data for more germplasms, transcriptome data from bulk, single-cell and spatial RNA-seq, epigenetic data from Hi-C, ATAC-seq or histone modification, etc. Therefore, future directions for SoyOmics mainly focus on continuous integration of these newly-generated omics data. In addition, artificial intelligence (AI)-based approaches for deep mining of these big data would provide valuable insights for a wide range of soybean studies, particularly for AI breeding in the era of big data. Towards this end, we would like to call for global collaborations to build SoyOmics as a valuable platform for the whole research community around the world.

## SUPPLEMENTAL INFORMATION

Supplemental information is available at Molecular Plant Online.

## AUTHOR CONTRIBUTIONS

Z.T., and Z.Z. conceived this project. Z.T., S.S., and Z.Z. supervised this work. Y.L., Y.S., X.L., and Y.Z. designed the framework of database and wrote the pipelines. S.L., and X.Y. revised germplasm information and phenotype records. X.L. and Y.Z. constructed the database. L.N. developed the pipeline used in VersionMap module. D.T. built up the easyGWAS, VersionMap and graph-based genome module. Y.L., Y.Z., X.N., S.Y., and Z.T. wrote the manuscript. Z.T., S.S., and Z.Z. revised the manuscript.

All authors read and approved the final manuscript.

## ACKNOWLEDGMENTS

We thank a number of SoyOmics users for their kind advice. No conflict of interest is declared.

## Materials and methods

### Gene and gene family functional annotation

The 27 *de novo* assembly soybean genomes were annotated by Pfam, InterPro, UniProt, and GO functional knowledgebases. We used InterProScan (version 5.52-86.0) (Jones et al., 2014) to annotate the protein sequences with Pfam and InterPro items. Then BLASTP results of protein sequence against the UniProt Swiss-Prot data were used for UniProt item annotation. Items matching the highest BLAST score, with identity ≥ 30% and length match portion ≥ 50% were treated as the UniProt annotation of genes. Furthermore, we used SANPANZ3, a stand-alone version of PANNZER2 (Toronen et al., 2018), to annotate proteins with GO item and functional description, by ppv over 0.4 as filtration. The KEGG pathway annotation was done with KofamScan (Aramaki et al., 2020).

To annotate the function of a gene family, we considered all the functional annotation items of its gene members and built an integrated method for gene group annotation. Suppose that one functional item is annotated to an *n*-sample homologous group with *m* (1 ≤ *m* ≤*n*) samples, we define that *N*_*i*_ is the total number of genes for any given sample *i* (*i* = 1, 2, 3…*n*) in the homologous group and *X*_*i*_ is the number of genes for the sample *i* that are annotated by the item. Thus, the annotation score (*S*) was formulated as:

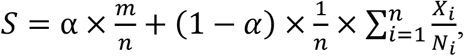

where α is the weight parameter for balance of the two measure aspects. In our study, α was set at 0.75. Finally, for each type of annotation base, such as Pfam, GO, UniProt, the item with the highest *S* value was picked as the annotation of the group.

For functional description of a homologous group, if any description text is the largest description and 1) is the unique; or 2) possesses over half of the genes and is 1.2 fold larger than the second one; or 3) possesses over thirty percent genes and is 2 fold larger than the second one, we determined the description text as the representative for the group.

### Modification of phenotype data

In SoyOmics, we collected 115 phenotypes, with 18 qualitative traits and 97 quantitative traits. To estimate the relationship between genotype and phenotype, firstly, we coded the 18 qualitative traits by numeric values (Supplemental Table 1). After that, 81 phenotypes collected from multi-areas or -years were modified by best linear unbiased prediction (BLUP) with R package “lme4”. Data passed through this modification were used for genotype-to-phenotype analysis.

### Selective test

Selective test was performed with the SNPs from the 2,898 soybean accessions. Tajima’s D (Tajima, 1989), reduce of diversity (ROD) (Xu et al., 2011), and F_ST_ (Weir and Cockerham, 1984) were calculated with 20 kb bin without overlapping. For Tajima’s D, the bottom 5% value was used as the significant threshold. For ROD and F_ST_, the top 5% value was chosen as the significant threshold.

### Variation imputation

To get the phased and no gap variation, we conducted imputation for the SNP and INDEL data of 2,898 soybeans. We split variants into the 5 Mb sections and used Beagle (v4.0) (Browning and Browning, 2007) to do the imputation, with parameters of phase-its=3 and impute-its=3. Then the separated variations were combined together by the chromosome and position rank.

### Variation effects estimation

In SoyOmics, we provided the Sorting Intolerant From Tolerant (SIFT) (Ng and Henikoff, 2001) values for SNPs to estimate their potential effects on the changing of amino acid. To do this analysis, the protein sequences from RefSeq and Swiss-Prot were used as target database. The SNP tagged by ‘non-synonymous SNV’, ‘stop-gain’ and ‘stop-loss’ were extracted as query variations. The results with Median Info within 2.75∼3.25 were treated as confident results and shown in SoyOmcis. If SIFT score was over 0.05, the substitution was tagged by Tolerated; otherwise, it was tagged by Damaging.

### Identification of MADS-box

Previous study identified a total of 157 *MADS-box* members in soybean (Gramzow and Theissen, 2013; Shu et al., 2013), with 131 having isoforms in the ZH13 genome. Pfam screening of ZH13 genome identified 184 genes that contained SRF-TF (PF00319) and/or K-box (PF01486) domain, which were the characteristic domains of *MADS-box*. Of the 184 genes, 130 were same as the reported genes. The genes with PF00319 or PF01486 was used for functional display as an example of *ExpPattern*.

### Generate chain file for genome position conversion

In SoyOmics, we provided the Genome position function between ZH13 and other soybean genomes. The chain files were generated with the UCSC liftOver pipeline (Kuhn et al., 2013). For each genome with ZH13, we generated two chain files by ZH13 as query or target. Therefore, users can do the bi-directional conversion between any genome and ZH13. The position conversion was done by CrossMap (Zhao et al., 2014) in SoyOmics.

---

**Supplemental Figure 1.**
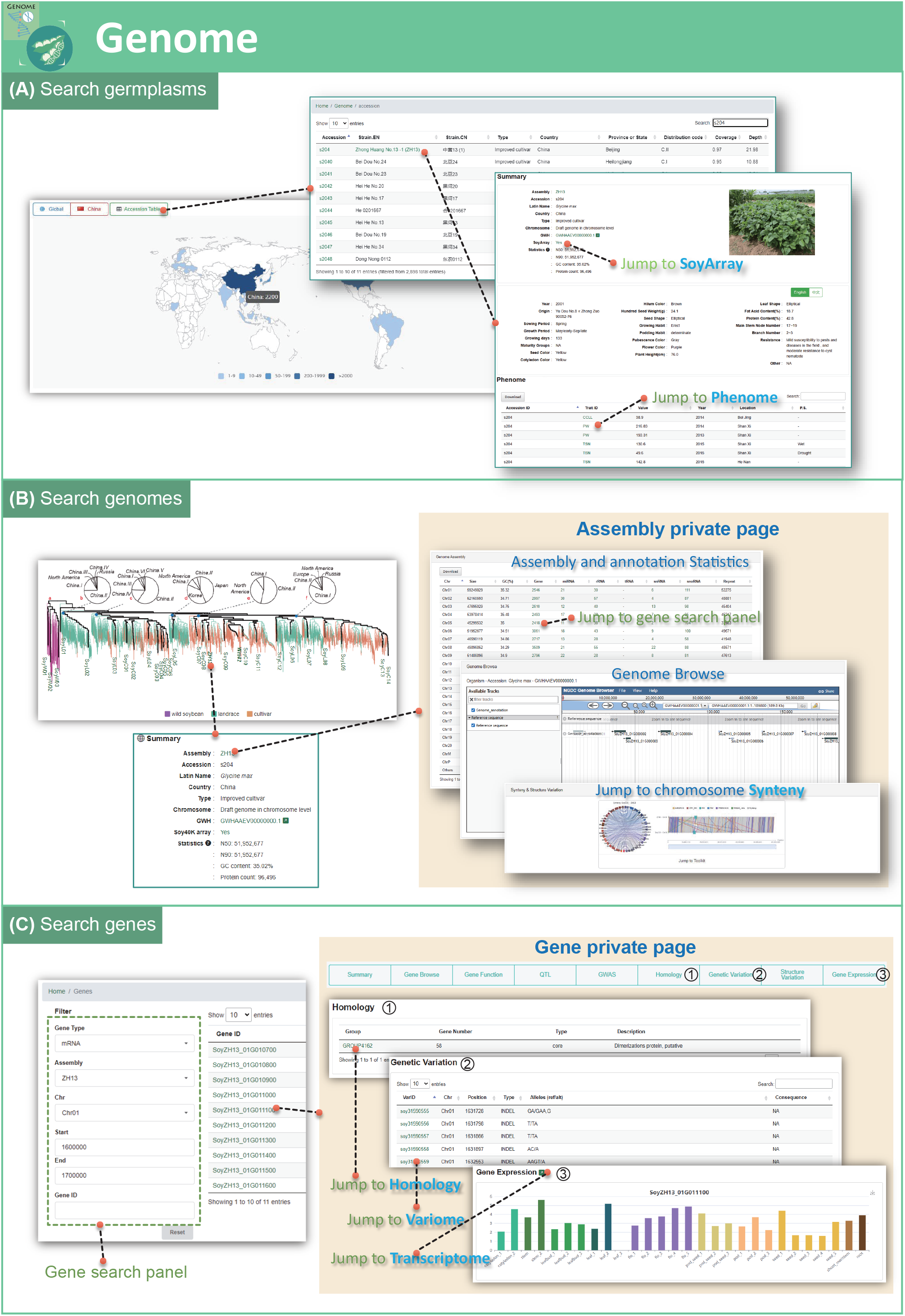
Introduction of Genome module. (A) Result of searching by germplasm. User can redirect to Phenome module and SoyArray tool from germplasm private page. (B) Result of searching by genome. Users can redirect to gene search panel and Synteny module from the assembly private page. (C) Result of searching by gene. Search panel affords setting of genome, gene type, genomic position and gene ID. Users can redirect to Variome, Homology and Transcriptome module from the gene private page.

**Supplemental Figure 2.**
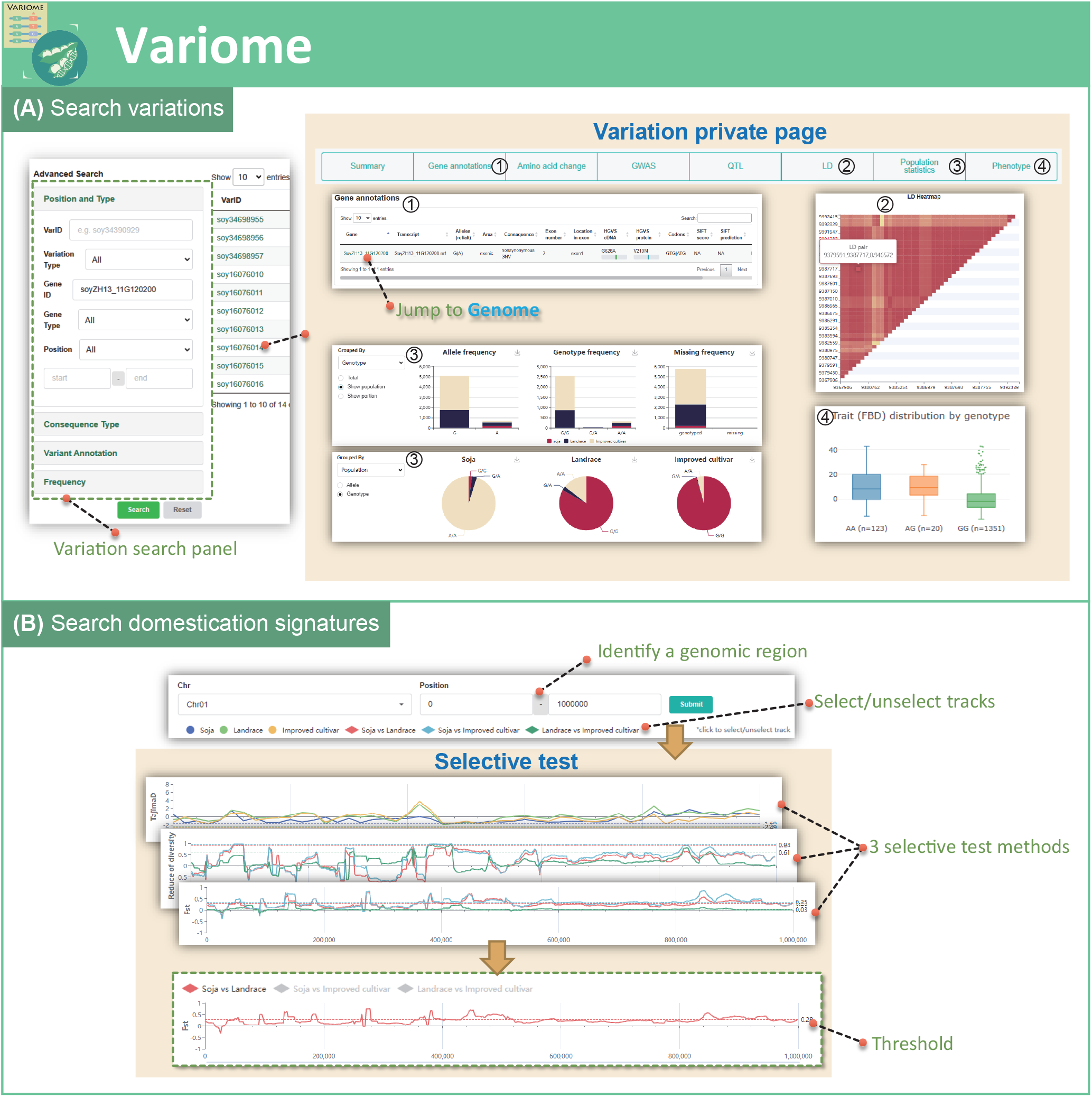
Introduction of Variome module. (A) Result of searching by variation. Search panel affords setting of genomic position, variation type, consequence on sequence, present knowledge and frequency. Users can redirect to gene private page and Phenome modules from variation private pate. (B) Result of searching by domestication signature. Track can be select/unselect to modify the visualization result.

**Supplemental Figure 3.**
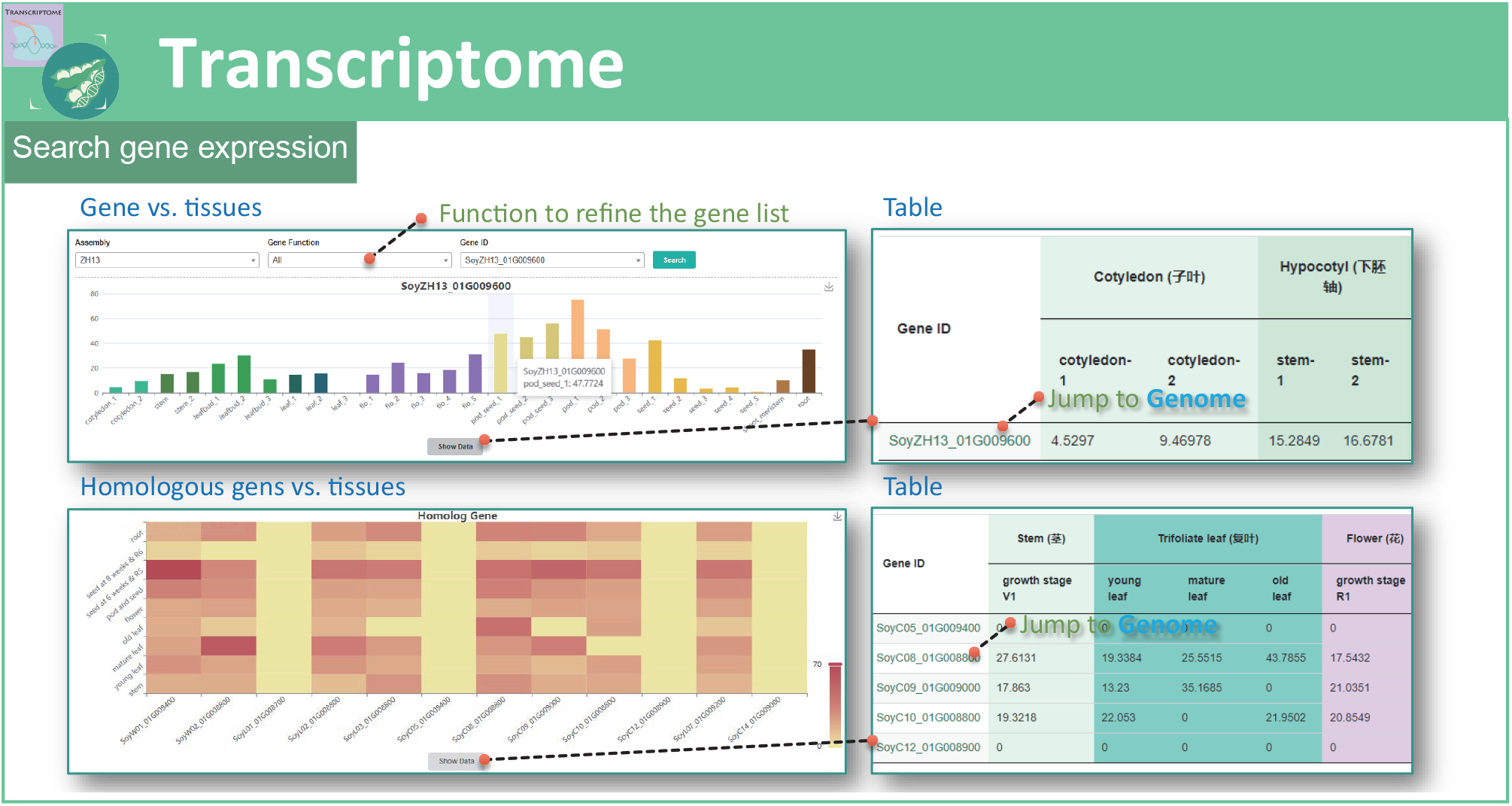
Introduction of Transcriptome module. This module shows expression of gene and their orthologous among tissues. Users can redirect to the gene private page for here. Expression value table can be hidden/appeared from here.

**Supplemental Figure 4.**
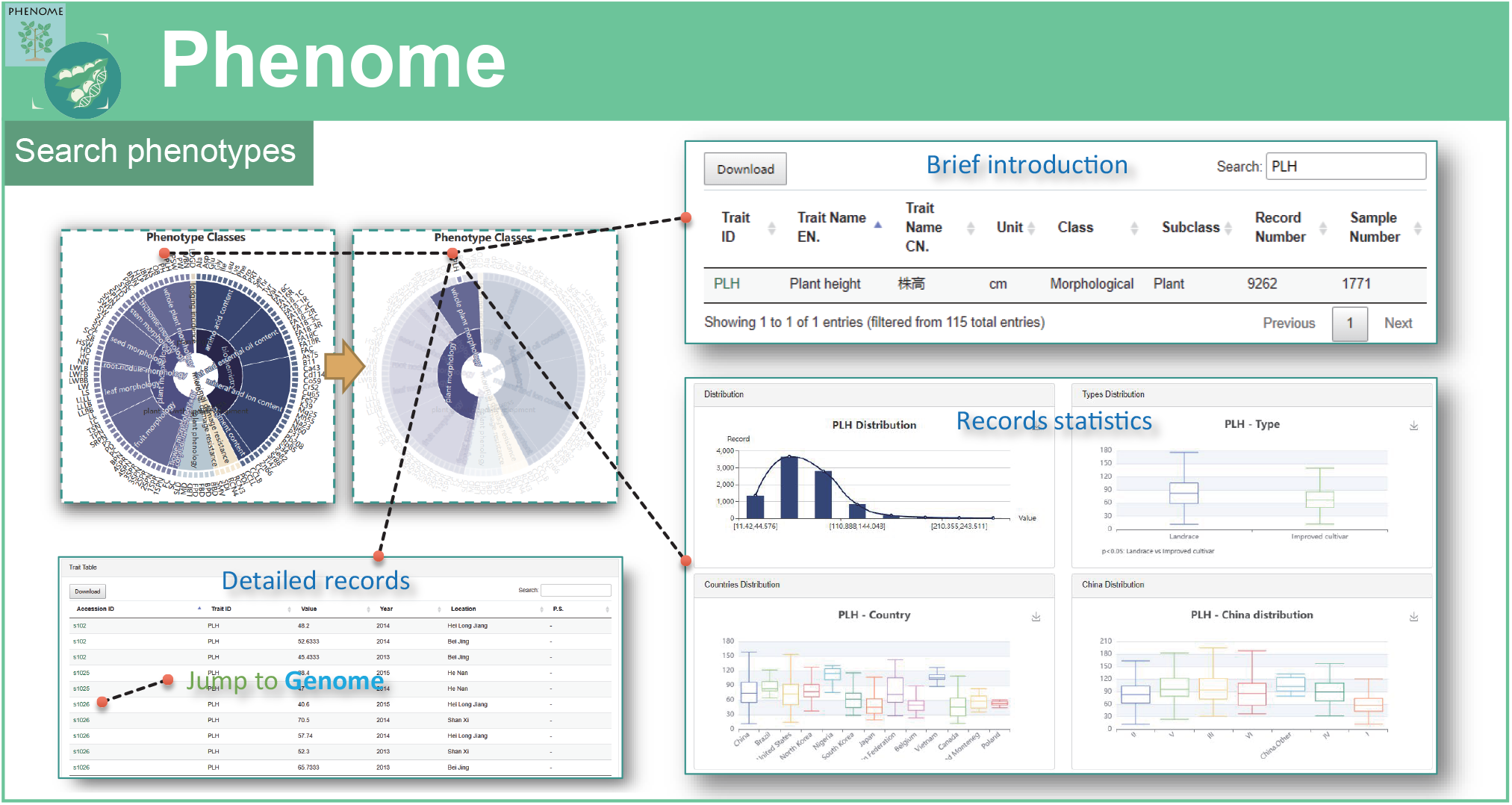
Introduction of Phenome module. This module shows introduction, records and statistics of phenotypes. Users can redirect to the germplasm private page from here.

**Supplemental Figure 5.**
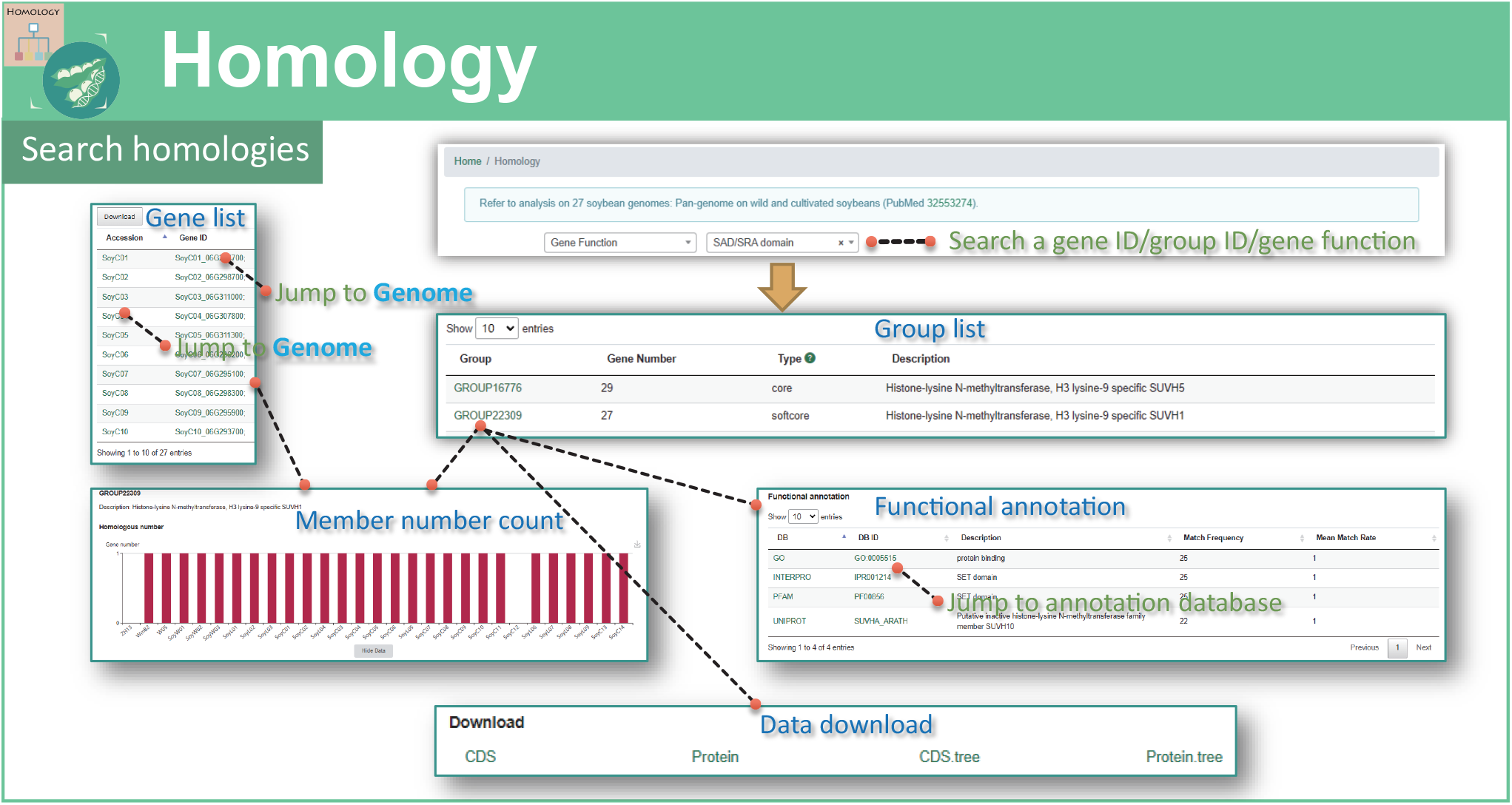
Introduction of Homology module. This module shows a homologous gene family with gene members among soybean genomes and the integrated annotation. Users can redirect to the gene private page here and download the CDS/protein sequences and phylogeny tree.

**Supplemental Figure 6.**
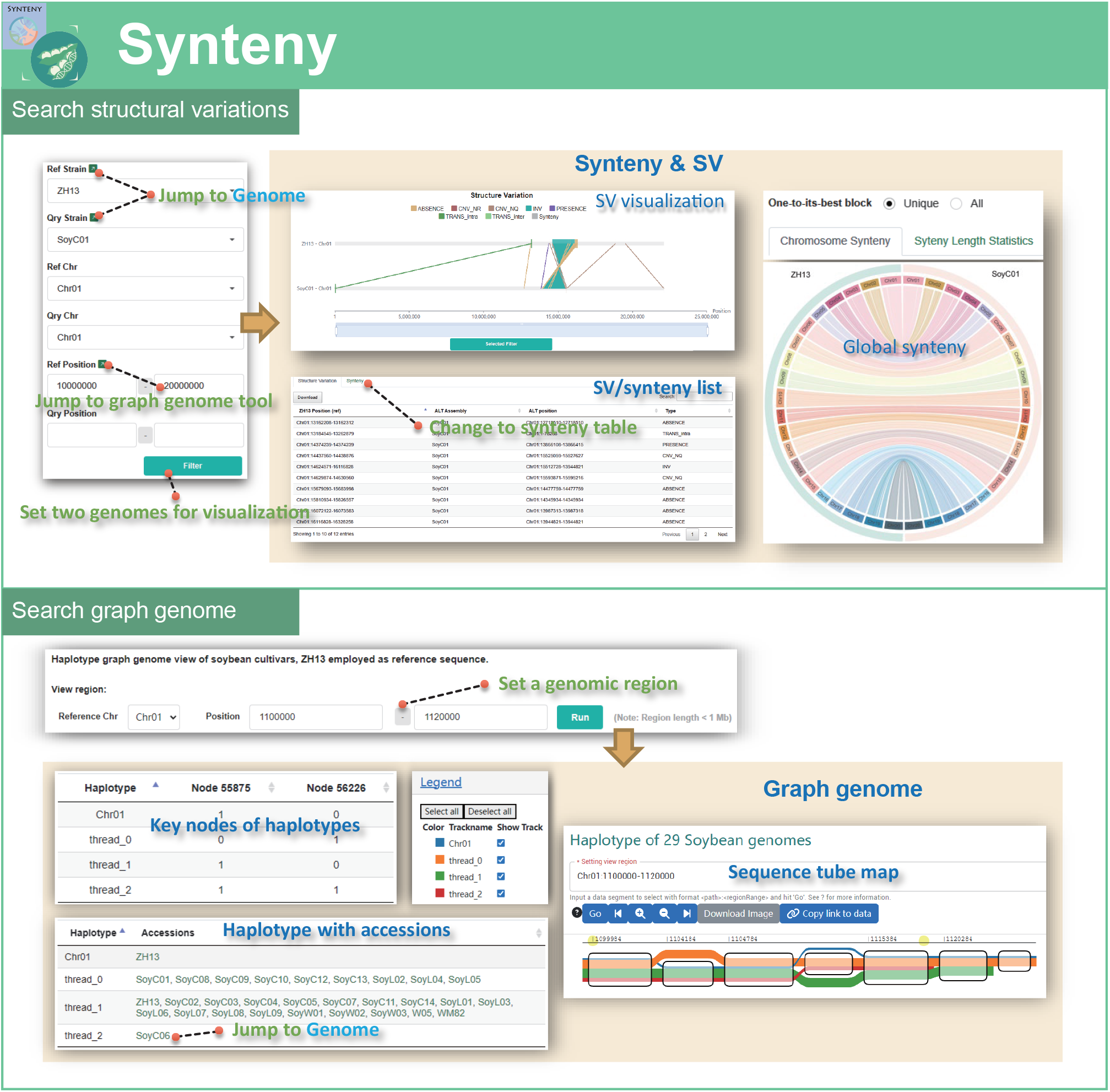
Introduction of Synteny module. (A) Chromosome synteny visualization and list of SVs between ZH13 and other soybean genomes. Users can redirect to any *de novo* assembled genome from here. (B) Graph genome visualization and haplotype identification according to nodes of graph.

**Supplemental Figure 7.**
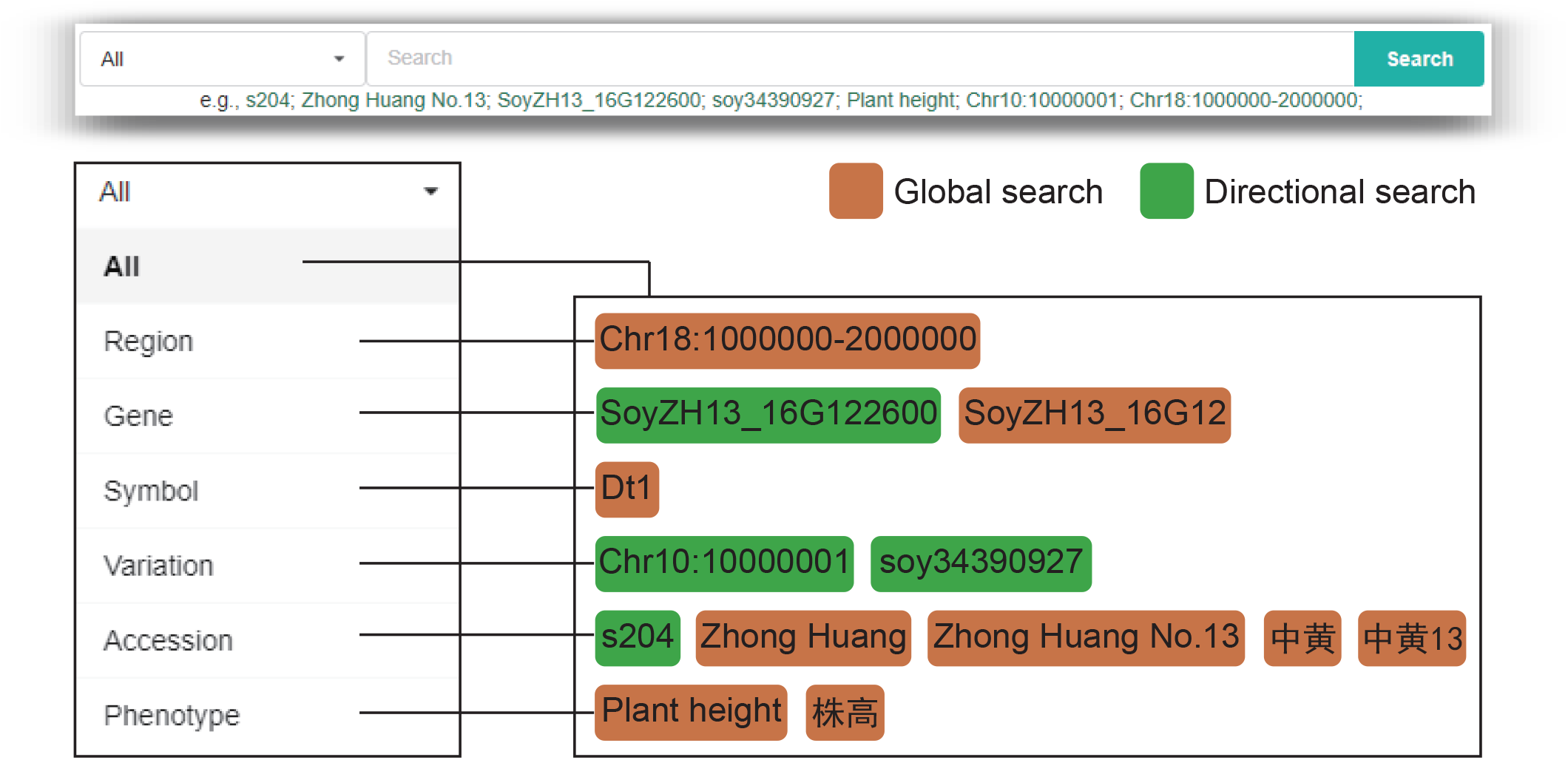
The comprehensive logic of searching bar. The global search provides all related result of the searched item in the SoyOmics. The directional search exactly jumps to the private page of the searched entity.

**Supplemental Figure 8.**
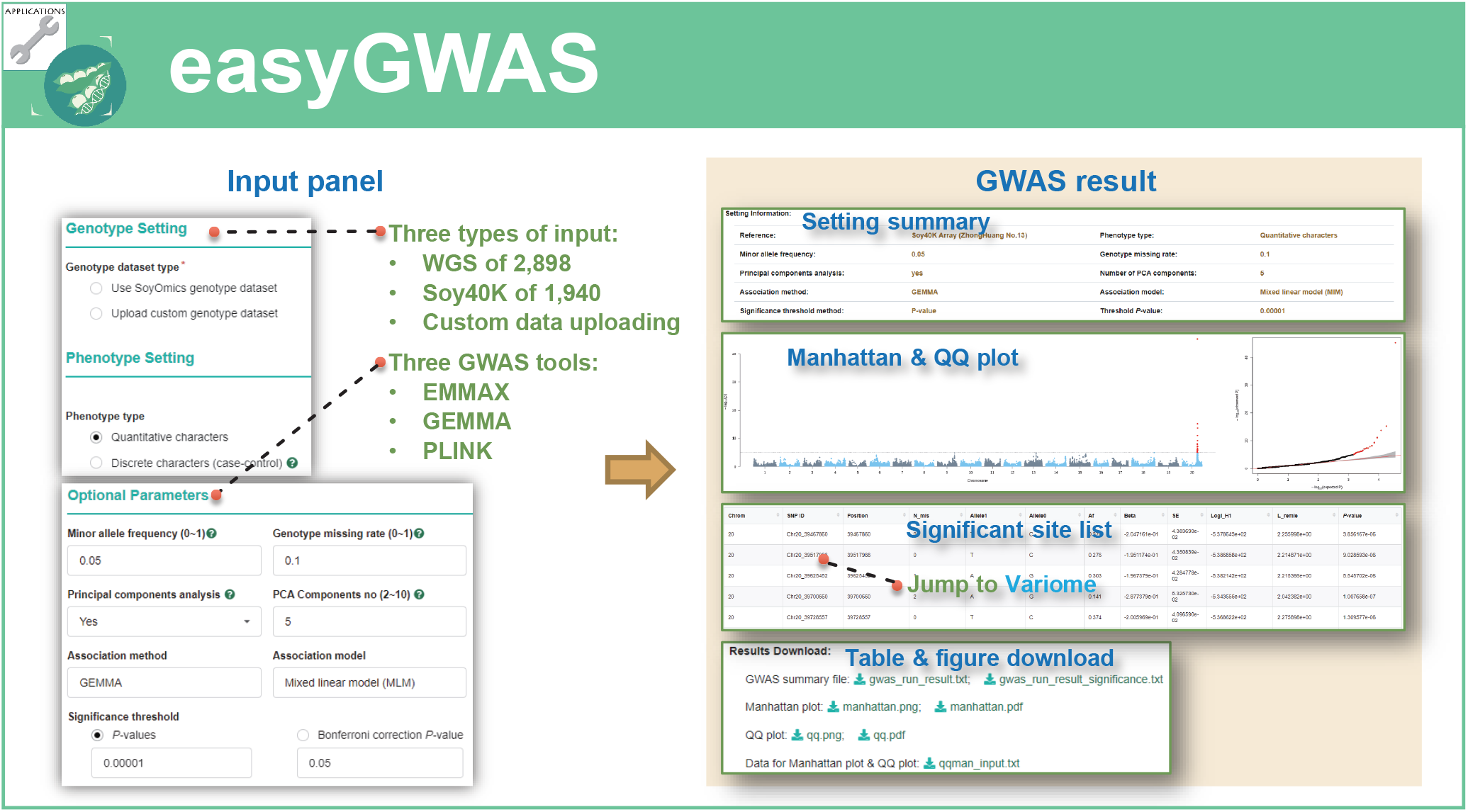
Guidance of *easyGWAS* with a case study of leaf width.

**Supplemental Figure 9.**
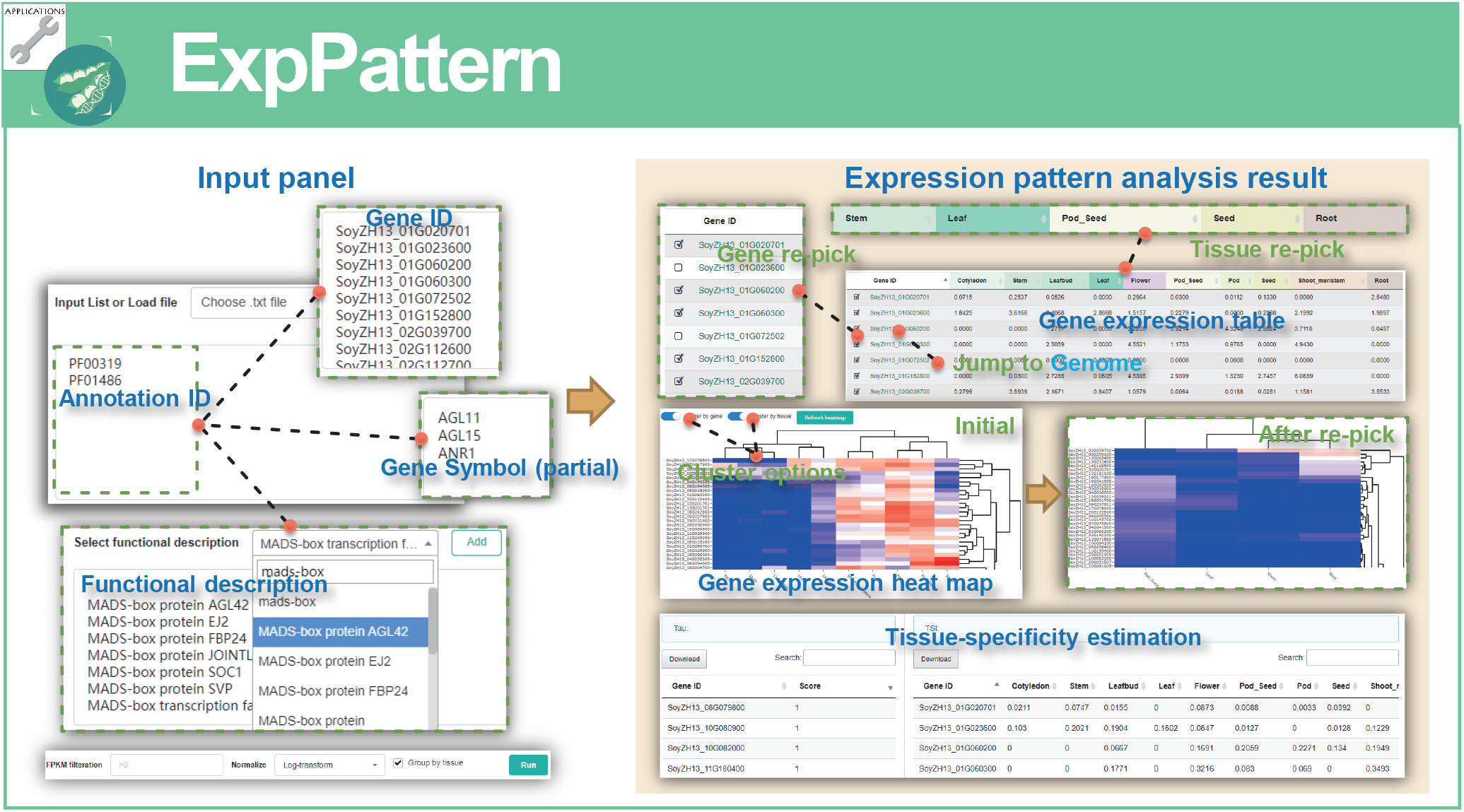
Guidance of *ExpPattern* with a case study of *MADS-box* family.

**Supplemental Figure 10.**
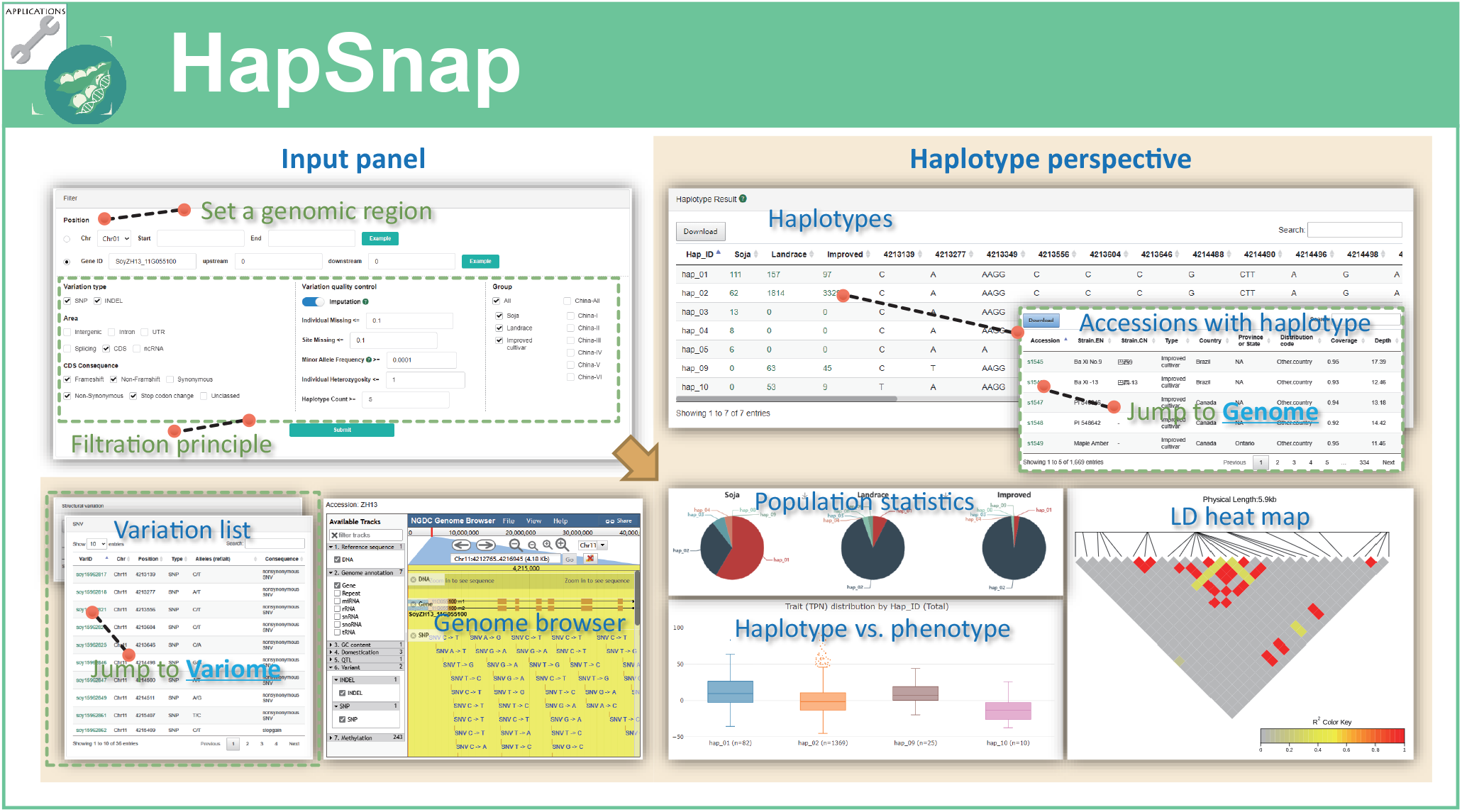
Guidance of *HapSnap* with a case study of *GmZEP*.

**Supplemental Figure 11.**
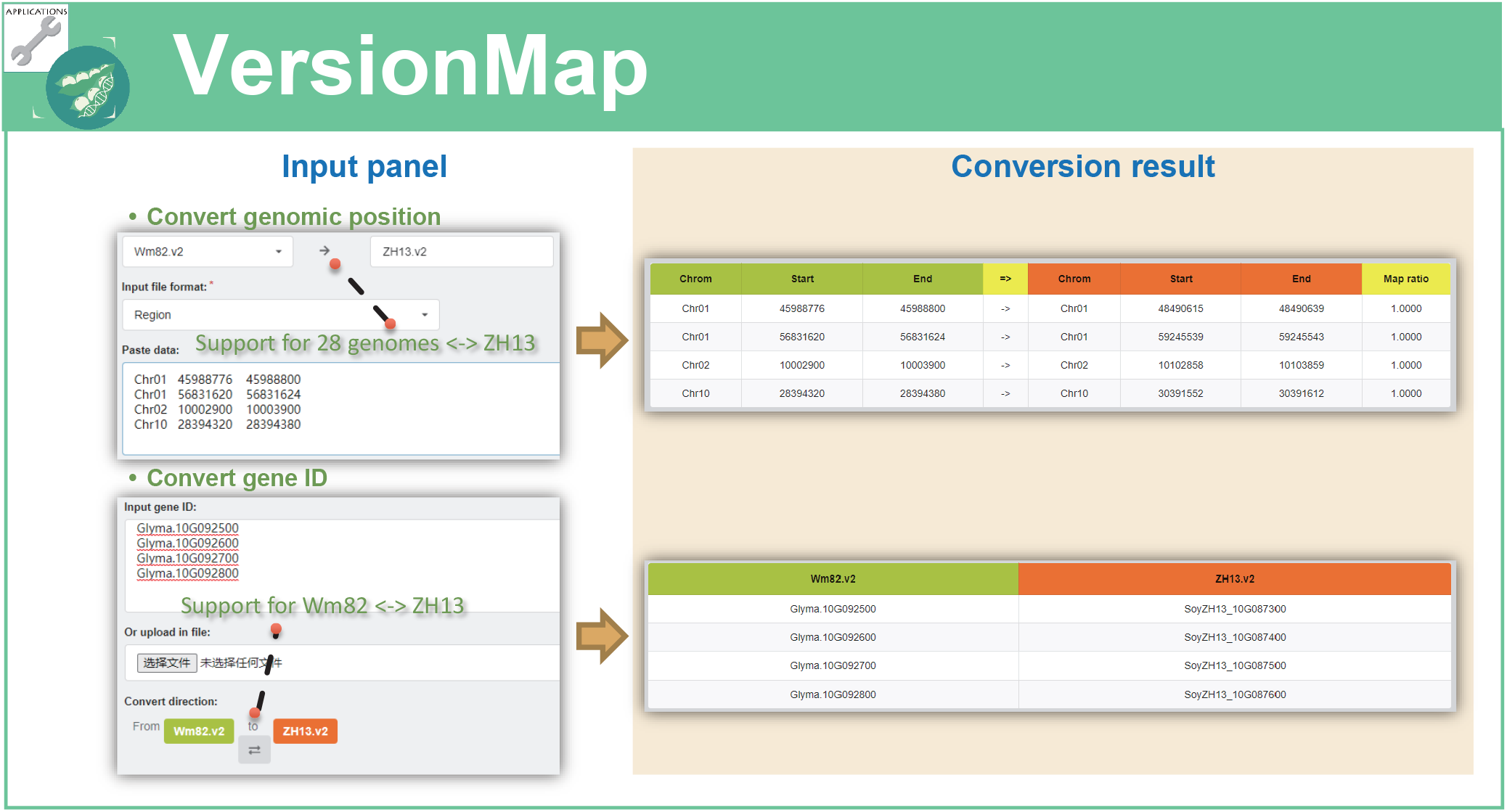
Guidance of *VersionMap* with examples of genomic regions and gene IDs.

**Supplemental Figure 12.**
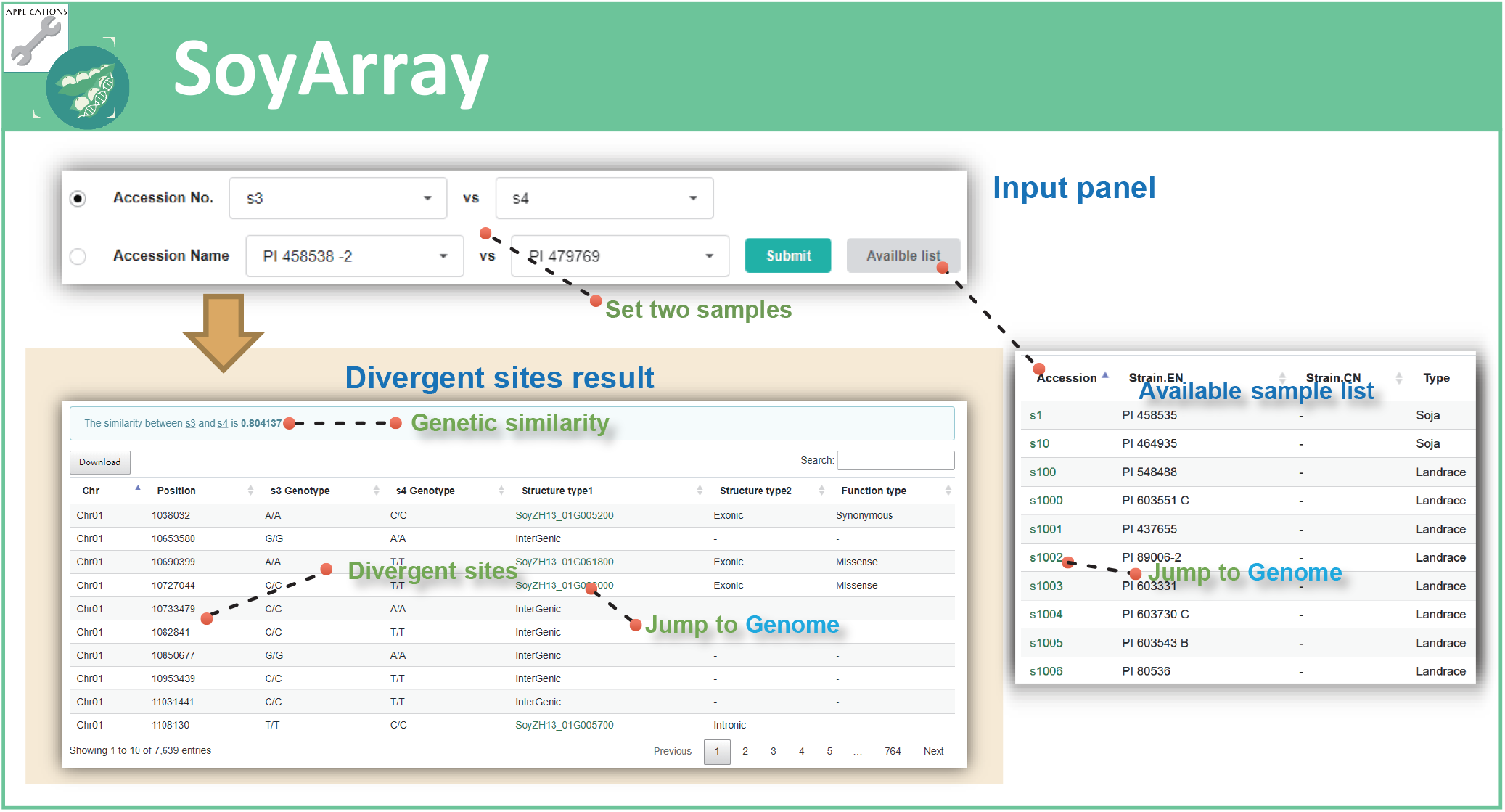
Guidance of *SoyArray* with comparison of two soybean accessions.

**Table S1.** Encoding of qualitative phenotypes.

| **Trait** | **Trait ID** | **Description** | **Code** |
| --- | --- | --- | --- |
| Resistance to cyst nematode 3 | RCN3 | Not | 1 |
| Resistance to cyst nematode 3 | RCN3 | Resistance | 2 |
| Resistance to cyst nematode 4 | RCN4 | Not | 1 |
| Resistance to cyst nematode 4 | RCN4 | Resistance | 2 |
| Flower color | FC | White | 1 |
| Flower color | FC | Purple | 2 |
| Leaf color | LC | Light Green | 1 |
| Leaf color | LC | Green | 2 |
| Leaf color | LC | Dark green | 3 |
| Leaf shape | LS | Lanceolate | 1 |
| Leaf shape | LS | Oval | 2 |
| Leaf shape | LS | Oval | 3 |
| Leaf shape | LS | Round | 4 |
| Branch shape | NS | Round | 1 |
| Branch shape | NS | Sector | 2 |
| Branch shape | NS | Single stem | 3 |
| Branch shape | NS | Fasciate | 4 |
| Podding habit | PH | Indeterminate | 1 |
| Podding habit | PH | Semi-determinate | 2 |
| Podding habit | PH | Determinate | 3 |
| Pod color | PC | Green | 1 |
| Pod color | PC | Yellow | 2 |
| Pod color | PC | Taupe | 3 |
| Pod color | PC | Brown | 4 |
| Pod color | PC | Dark brown | 5 |
| Pod color | PC | Black | 6 |
| Pubescence color | PBC | Gray | 1 |
| Pubescence color | PBC | Brown | 2 |
| Seed shape | SS | Round | 1 |
| Seed shape | SS | Oblate | 2 |
| Seed shape | SS | Oval | 3 |
| Seed shape | SS | Long oval | 4 |
| Seed shape | SS | Reniform | 5 |
| Seed shape | SS | Other | 6 |
| Seed brightness | SB | No brightness | 1 |
| Seed brightness | SB | Slight brightness | 2 |
| Seed brightness | SB | Strong brightness | 3 |
| Seed coat color | SCC | Yellow | 1 |
| Seed coat color | SCC | Green | 2 |
| Seed coat color | SCC | Black | 3 |
| Seed coat color | SCC | Brown | 4 |
| Seed coat color | SCC | Mixed | 5 |
| Seed coat color | SCC | Other | 6 |
| Hilum diffusion | HD | Not diffused | 1 |
| Hilum diffusion | HD | Less Diffused | 2 |
| Hilum diffusion | HD | Diffused | 3 |
| Hilum color | HC | Yellow | 1 |
| Hilum color | HC | Light brown | 2 |
| Hilum color | HC | Brown | 3 |
| Hilum color | HC | Black | 4 |
| Hilum color | HC | Dark brown | 5 |
| Hilum color | HC | Blue | 6 |
| Hilum color | HC | Light black | 7 |
| Hilum color | HC | Other | 8 |
| Seedling color | SC | Purple | 1 |
| Seedling color | SC | Green | 2 |
| Seedling color | SC | White | 3 |
| Seedling color | SC | Brown | 4 |
| Stem shape | STS | Normal | 1 |
| Stem shape | STS | Fasciate | 2 |
| Stem shape | STS | Curving | 3 |
| Stem shape | STS | Other | 4 |
| Pod shattering | PS | Not shattering | 1 |
| Pod shattering | PS | Shattering | 2 |
| Soybean mosaic virus | SMV | Not | 1 |
| Soybean mosaic virus | SMV | Resistance | 2 |

